# Functional redundancy in chicken ANP32A mediates species-specific support of avian influenza virus polymerase

**DOI:** 10.1101/2025.03.25.645163

**Authors:** Liuke Sun, Yuxing Qu, Xue-Feng Wang, Mengmeng Yu, Xiaojun Wang

## Abstract

Host restriction of avian influenza virus (AIV) polymerase in human cells is driven by species-specific differences in ANP32A/B proteins. While AIV polymerase relies on ANP32A/B containing a 33-amino-acid insert unique to avian species, the structural and/or mechanistic basis for this requirement remains poorly characterized. Here, we demonstrate that chicken ANP32A (chANP32A) displays three functional determinants enabling its species-specific support of AIV polymerase: (1) a SUMO-interacting motif (SIM), (2) SUMOylation at residues K68/K153, and (3) a 28-amino-acid segment within the avian-specific insertion. These determinants function synergistically and redundantly, requiring at least two for optimal activity, to enhance interactions between AIV viral ribonucleoprotein (vRNP) and chANP32A, thereby promoting AIV vRNP assembly. In contrast, human ANP32A/B—which lack the other two determinants and rely solely on SUMOylation—exhibit a limited capacity to support AIV polymerase activity. Our findings unveil a cooperative mechanism where SUMO-dependent processes and structural motifs in chANP32A enforce species-specific adaptation of AIV polymerase, shedding light on how ANP32A/B governs host restriction of AIV polymerase.

## Introduction

Aquatic birds serve as the main natural reservoir for avian influenza viruses (AIVs) ^1,2^. These viruses can transmit across species barriers to infect humans, either directly or via intermediate hosts such as pigs ^3–5^. Although spillover events from avian hosts to mammals occur frequently, innate host-specific barriers restrict sustained transmission ^3–6^. The first major barrier is the low binding affinity of the AIV hemagglutinin (HA) protein for human receptors^7^, which hinders viral entry into human cells. The second key barrier stems from the poor activity of the AIV RNA-dependent RNA polymerase (vPol) in mammalian cells, inhibiting efficient viral genome replication and transcription^6^. Despite these obstacles, certain AIVs can occasionally overcome these barriers, leading to successful cross-species transmission and posing a significant threat to human populations ^8–15^.

ANP32 proteins, including paralogues ANP32A, ANP32B, and ANP32E, are characterized by an N-terminal leucine-rich repeat (LRR) domain and a C-terminal low-complexity acidic region (LCAR), separated by a central domain. These proteins play diverse roles in histone chaperoning, transcriptional regulation, nuclear export, and apoptosis ^16–18^. Recent work has identified ANP32A/B as critical cofactors for influenza A, B, and C virus polymerases, with species-specific variations in these proteins dictating viral host range ^19–24^. In chickens, an exon duplication introduces a 33-amino-acid insert within the central domain of ANP32A, generating a longer isoform that robustly supports AIV polymerase activity^19^. In contrast, mammals express truncated ANP32A/B isoforms lacking this insertion, which inefficiently support AIV polymerase, forming a major barrier to avian virus replication in mammalian cells^19,25,26^. Functional redundancy among paralogues further differs between hosts: human ANP32A and ANP32B (huANP32A/B) act redundantly to sustain influenza A polymerase activity, whereas chicken ANP32B is evolutionarily inactive, leaving chANP32A as the sole functional cofactor in chickens^20–22^. Neither chicken nor mammalian ANP32E contributes to influenza A polymerase function^23^. The restriction of AIV polymerase activity in mammalian cells can be circumvented by either overexpressing chANP32A or acquiring adaptive mutations (e.g., PB2-E627K), which frequently emerge during AIV evolution in mammalian hosts ^19,20,27^. Despite strong genetic evidence linking ANP32A/B divergence to AIV’s host range limitations, the precise structural or mechanistic basis through which chANP32A selectively enhances avian polymerase activity remains unresolved.

SUMOylation is an essential post-translational modification involving the covalent attachment of SUMO molecules to lysine residues on target proteins^28,29^. This process is mediated by a cascade of enzymatic reactions involving the activation enzyme E1 (SAE1/UBA2), the conjugation enzyme E2 (Ubc9) and E3 SUMO ligases. In mammals, three SUMO isoforms (SUMO1-3) contribute to SUMOylation processes. The modification typically occurs at a conserved consensus motif (ҨKxD/E), where Ҩ represents a bulky hydrophobic group^30,31^. SUMO-dependent functions can also be mediated by non-covalent interactions between SUMO and substrates via SUMO-interacting motifs (SIMs), facilitated by the SUMO-SIM binding mechanism^32,33^. In chANP32A, a functional SIM embedded within its unique 33-amino-acid insertion is essential for robust support of AIV polymerase activity^27,34,35^. Intriguingly, the chANP32A-X2 isoform, a naturally occurring variant lacking an intact SIM, retains strong polymerase-supporting capacity, suggesting compensatory mechanisms or unidentified functional elements within chANP32A. Our recent work demonstrated that huANP32A/B undergo SUMOylation—a process exploited by the AIV NS2 protein to subvert host restriction barriers ^36^. Nevertheless, key mechanistic questions persist: (1) Does the SIM in chANP32A promote its own SUMOylation? (2) What functional role does SUMO modification play in chANP32A’s ability to enhance AIV polymerase activity?

In this study, we elucidate three redundant determinants in chANP32A —a SIM, SUMOylation, and a 28-amino-acid (28-aa) insert—that collectively govern its ability to support AIV polymerase activity. Strikingly, any two of these elements are sufficient for robust support. The SIM and SUMOylation operate via SUMO-dependent mechanisms and can functionally compensate for one another. While the 28-aa insert functions independently of direct SUMOylation, it requires SUMO-dependent activity—either through the SIM or SUMOylation—to exert its effect. Notably, the AIV polymerase-supporting activities of the 28-aa insert and SIM can be engineered into huANP32A, but only the SIM-mediated activity transfers to huANP32B. Since both human orthologs lack the 28-aa insert and SIM—relying solely on SUMOylation—they fail to efficiently sustain AIV polymerase activity. Mechanistically, three determinants enable chANP32A to enhance AIV polymerase function by strengthening interactions between AIV vRNP and chANP32A, and promoting AIV vRNP assembly. These insights resolve longstanding questions about the molecular basis of ANP32A/B-mediated host restriction of AIV polymerase, underscoring the combinatorial role of structural and post-translational elements in viral adaptation.

## Results

### Chicken ANP32A harbors functional determinants beyond the SIM for supporting AIV polymerase activity

The avian-specific 33-amino-acid insert of chANP32A contains a SIM (residues 176-VLSLV-180) essential for supporting AIV polymerase activity^37^. Three naturally occurring chANP32A isoforms exhibit variations in this insert^27,34,35^: the wild-type isoform (chANP32A-X1) retains the full 33-amino-acid sequence, chANP32A-X2 lacks four residues within the SIM (retaining V180 and a truncated 28-aa insert), and chANP32A-X3 mirrors mammalian isoforms by lacking the insert entirely (Fig. 1A). To further dissect the SIM’s functional role, we therefore took advantage of the three different chANP32A isoforms that exist in nature by using the previously described *ANP32A&B&E*-triple-knockout HEK293T cells (HEK293T-TKO)^23^ to establish an ANP32-dependent mini-replicon polymerase reporter system through complementing polymerase function with the expression of different ANP32 proteins. Consistent with prior studies^34,35^, chANP32A-X1 exhibited the strongest support for H9N2 and H7N9 (PB2-627E) polymerase activity. Notably, chANP32A-X2, despite its disrupted SIM, retained substantial (though reduced) capacity to facilitate AIV polymerase activity, whereas chANP32A-X3 and huANP32A showed minimal support (Fig.1B and C). To correlate these findings with viral replication, we used the previously described ANP32A&B&E triple-knockout MDCK cells (MDCK-TKO)^36^ to generate different cell lines stably expressing Flag-tagged chANP32A isoforms. Western blot analysis confirmed equivalent expression (Fig.1D). Multi-cycle replication assays of the H9N2 virus revealed that chANP32A-X1 produced the highest viral titers, followed by chANP32A-X2, with chANP32A-X3 exhibiting the lowest (Fig.1E).

**Figure 1.**
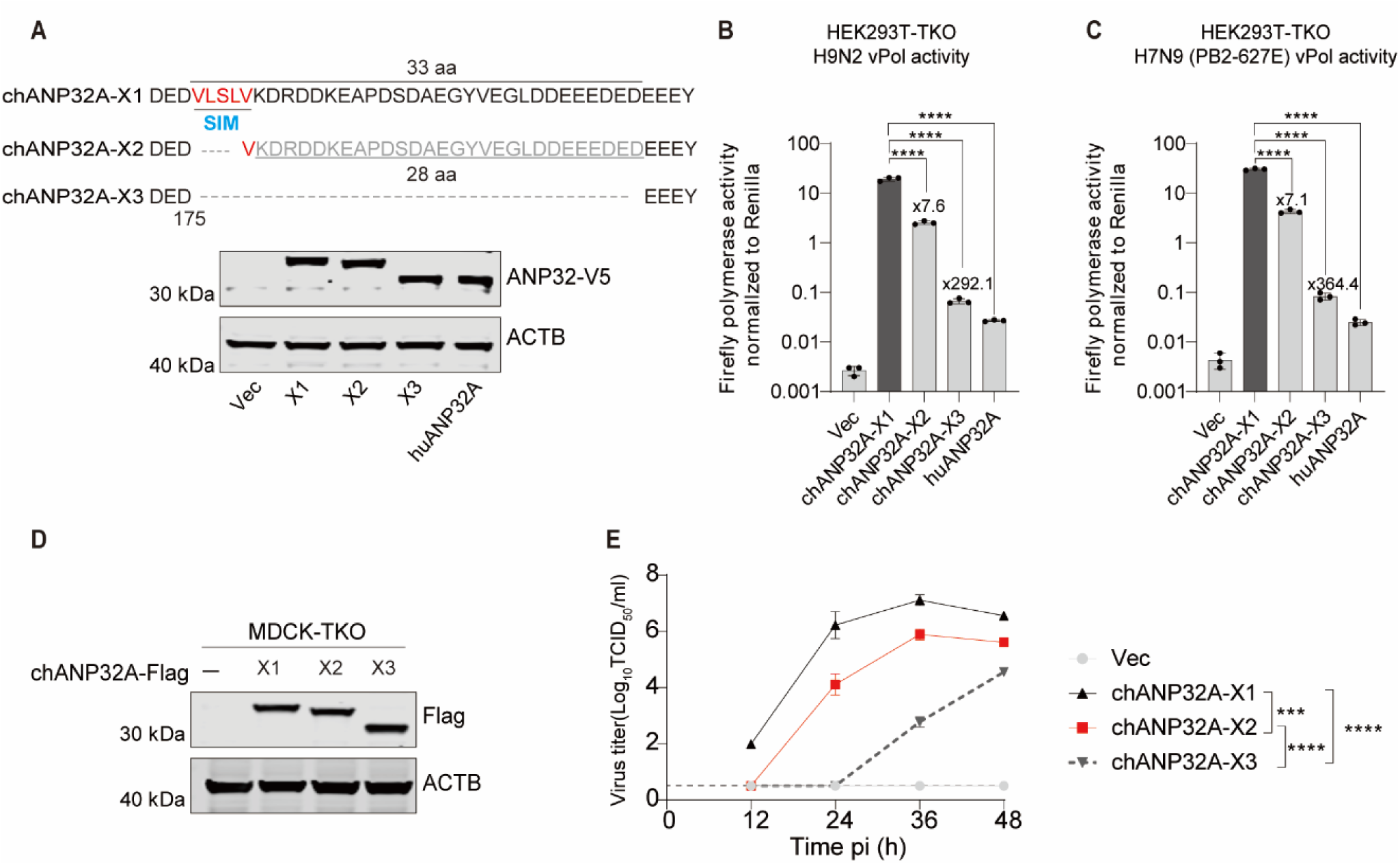
Chicken ANP32A-X2 lacks intact SIM but retains high ability to support AIV polymerase activity. **(A)** Sequence alignment of three chANP32A isoforms is shown. Western blots demonstrate comparable expression levels of various V5-tagged ANP32A constructs in HEK293T-TKO cells. (**B** and **C**) Polymerase reconstitution assay in HEK293T-TKO cells comparing the effect of each ANP32A-V5 construct on polymerase activity of H9N2 (B) and H7N9 (PB2-627E) (C). Statistical analysis and fold changes relative to X1 are presented. (**D**) Immunoblotting analysis of MDCK-TKO cells stably reconstituted with the indicated chANP32A-Flag constructs or an empty vector. (**E**) Replication kinetics of the avian H9N2 virus in control MDCK-TKO cells or those stably expressing Flag-tagged X1, X2, or X3 (MOI = 0.001). Viral titers were determined at the indicated time points. In (B), (C), and (E), error bars represent mean ± SD from *n* = 3 independent biological replicates. Statistical significance was determined by one-way ANOVA with Dunnett’s multiple comparisons (B and C) or two-way ANOVA (E) (NS, not significant; \*\*\**p* < 0.001; \*\*\*\**p* < 0.0001).

Taken together, these data suggest that the SIM is essential but not solely responsible for chANP32A’s ability to support AIV polymerase activity. The residual functionality of chANP32A-X2—despite SIM truncation—implies the existence of additional functional elements within the 29-amino-acid insert or other chANP32A domains that partially compensate for SIM disruption.

### Chicken ANP32A is a SUMOylation substrate

SUMO-dependent functions are mediated either by non-covalent SIM-SUMO interactions or covalent SUMOylation^28,29^. We hypothesized that chANP32A-X2’s retained ability to support AIV polymerase activity—despite lacking an intact SIM—might involve compensatory SUMOylation. To test this, we first assessed SUMOylation of full-length chANP32A (X1). HEK293T cells were co-transfected with chANP32A-HA, Myc-Ubc9, and His-tagged wild-type SUMO1/2/3 or non-conjugatable mutants (SUMO1m/2m/3m, with C-terminal glycine-to-alanine substitutions). SUMOylated proteins were enriched via Ni^2+^-NTA beads pulldown under denaturing conditions. Immunoblotting with anti-HA antibody revealed three distinct bands (>50 kDa) for chANP32A co-expressed with wild-type SUMO1/2/3, but not with SUMO mutants (Fig. 2A). Given chANP32A-HA’s molecular weight (∼33 kDa) and SUMO’s size (∼15–20 kDa), these results indicate poly-SUMOylation (likely tri-SUMO chains) on chANP32A. Additionally, both the SUMO1 conjugation process and chANP32A SUMOylation were markedly reduced by the SUMOylation inhibitor TAK-981 (Fig. 2B).

**Figure 2.**
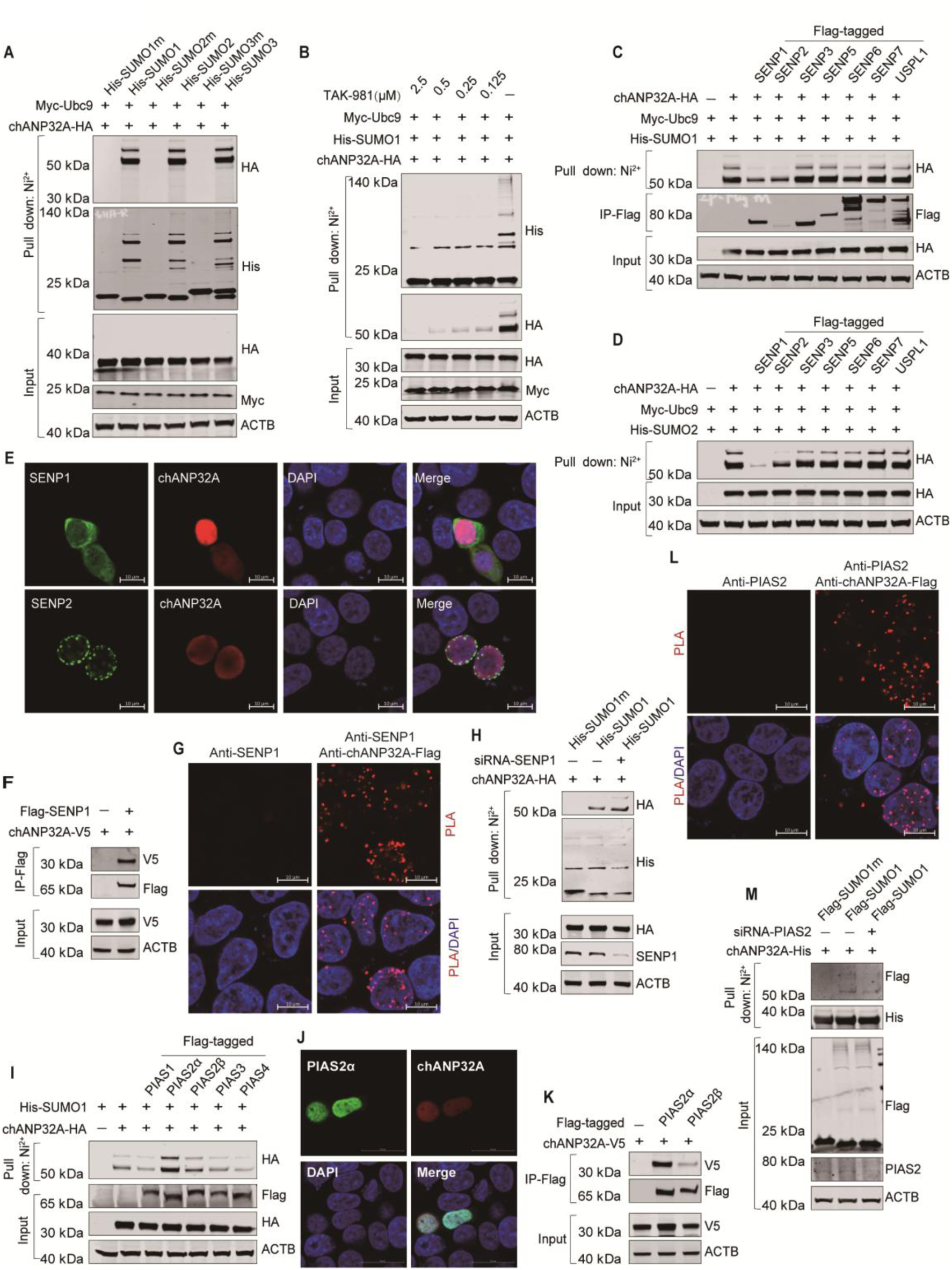
SUMOylation of chANP32A is induced by E3 SUMO ligase PIAS2α and reversed by SENP1. **(A)** Ni^2+^-NTA beads affinity pull-down assay demonstrating the SUMO modification of chANP32A with SUMO1, SUMO2, or SUMO3 in HEK293T cells. (**B**) Ni^2+^-NTA beads affinity pull-down assay results showing that TAK-981 treatment abolished chANP32A SUMOylation in HEK293T cells. HEK293T cells were transfected with the indicated plasmids. Twenty-four hours post-transfection, TAK-981 was added and incubated for an additional 24 hours prior to cell collection for the SUMOylation assay (**C, D**) Ni^2+^-NTA beads affinity pull-down assay revealing that overexpression of SENP1 or SENP2 inhibited chANP32A SUMOylation mediated by SUMO1 (C) or SUMO2 (D) in HEK293T cells. (**E**) Immunofluorescence staining analysis of co-localization between chANP32A-V5 and Flag-SENP1 or Flag-SENP2. Scale bar: 10 μm. (**F**) Co-IP experiments demonstrating the interaction between SENP1 and chANP32A in HEK293T cells. (**G**) Proximity ligation assay (PLA) confirming the interaction between endogenous SENP1 and chANP32A-Flag in HEK293T cells stably expressing chANP32A-Flag. Scale bar: 10 μm. (**H**) Ni^2+^-NTA beads affinity pull-down assay results showing enhanced chANP32A SUMOylation upon knockdown of endogenous SENP1 in HEK293T cells. (**I**) Ni^2+^-NTA beads affinity pull-down assay indicating that chANP32A SUMOylation was enhanced by overexpression of PIAS2α in HEK293T cells. (**J**) Immunofluorescence staining analysis of co-localization between chANP32A-V5 and Flag-PIAS2α. Scale bar: 20 μm. (**K**) Co-IP experiments confirming the interaction between PIAS2α and chANP32A in HEK293T cells. (**L**) PLA confirming the interaction between endogenous PIAS2 and chANP32A-Flag in HEK293T cells stably expressing chANP32A-Flag. Scale bar: 10 μm. (**M**) Ni^2+^-NTA beads affinity pull-down assay results showing chANP32A SUMOylation in HEK293T cells was notably reduced upon knockdown of PIAS2 in HEK293T cells. All the western blot experiments were independently repeated three times with consistent results.

In mammals, sentrin-specific proteases (SENPs) are the major enzymes that catalyze deSUMOylation^38^. To identify SENPs regulating chANP32A SUMOylation, we overexpressed nuclear SENPs (SENP1-3, SENP5-7, USPL1) in HEK293T cells. SENP1 and SENP2 overexpression considerably reduced chANP32A SUMOylation (SUMO1/SUMO2; Figs. 2C and D). Immunofluorescence revealed that SENP2 mainly localized to the nuclear envelope, whereas SENP1 exhibited both cytoplasmic and nuclear distribution. Notably, SENP1 displayed stronger nuclear co-localization with chANP32A compared to SENP2 (Fig. 2E). Therefore, we focused on addressing the role of SENP1 in chANP32A deSUMOylation. Co-immunoprecipitation (IP) confirmed direct interaction between Flag-SENP1 and chANP32A-V5 (Fig. 2F). Proximity ligation assays (PLA) in HEK293T cells stably expressing chANP32A-Flag validated endogenous SENP1-chANP32A interactions (Fig. 2G). Critically, SENP1 knockdown substantially enhanced chANP32A SUMOylation (Fig. 2H), establishing SENP1 as the dominant deSUMOylase for chANP32A.

The PIAS (protein inhibitor of activated Stat) family proteins, which include at least five members—PIAS1, PIAS3, PIAS4 (PIASy), and the α and β splice variants of PIAS2 (PIASx)—are the major SUMO E3 ligases. To identify chANP32A’s E3 SUMO ligase, each of the E3 ligases PIAS1–4 and His-SUMO1 were co-transfected with chANP32A-HA into HEK293T cells and chANP32A SUMOylation was assessed by performing SUMOylation assays. PIAS2α overexpression robustly enhanced chANP32A SUMOylation (Fig. 2I). Nuclear co-localization of PIAS2α with chANP32A was confirmed by immunofluorescence (Fig. 2J), and Co-IP demonstrated their direct interaction (Fig. 2K). PLA further validated endogenous PIAS2-chANP32A interaction in HEK293T cells stably expressing chANP32A-Flag (Fig. 2L). PIAS2 knockdown markedly reduced chANP32A SUMOylation (Fig. 2M), confirming its role as the primary SUMO E3 ligase.

Taken together, these findings demonstrate that chANP32A undergoes SUMOylation mediated by PIAS2α and reversed by SENP1.

### SUMOylation is a critical functional determinant in chANP32A-X1 and chANP32A-X2

To investigate whether SUMOylation compensates for the loss of the SIM in chANP32A-X2, we first confirmed its SUMO modification status. SUMOylation assays in HEK293T cells revealed that chANP32A-X2 and chANP32A-X3 exhibited SUMOylation patterns similar to chANP32A-X1 (Fig. 3A). We obtained consistent results without overexpression of Ubc9 (Fig. S1). Interestingly, we noted that levels of the second SUMOylation band in chANP32A-X2 were slightly higher than in chANP32A-X1, implying that the loss of the SIM does not reduce chANP32A SUMOylation, but rather promotes it To assess the functional importance of SUMOylation, we generated lysine-free mutants (chANP32A-X1-K0 and X2-K0) by replacing all lysines with arginines, thereby abolishing SUMOylation (Fig 3B). Viral polymerase activity assays in HEK293T-TKO cells revealed that that both X1-K0 (SUMOylation-deficient) and X2 (SIM-deficient) showed significantly reduced AIV polymerase activity compared to wild-type X1 (Fig 3C). Notably, we noticed that loss of either SIM (X2) or SUMOylation (X1-K0) alone partially impaired activity, whereas simultaneous loss of both SIM and SUMOylation nearly abolished activity, highlighting their functional redundancy. Multi-cycle replication assays in MDCK-TKO cells transiently expressing chANP32A mutants recapitulated these findings: wild-type X1 supported robust H9N2 replication, whereas X1-K0 and X2 displayed attenuated replication efficiency, and X2-K0 completely failed to sustain viral replication (Figs. 3D and E). These results collectively demonstrate that chANP32A employs compensatory SUMO-dependent mechanisms—covalent SUMOylation and non-covalent SIM interactions—to maintain its critical support of AIV polymerase activity.

**Figure 3.**
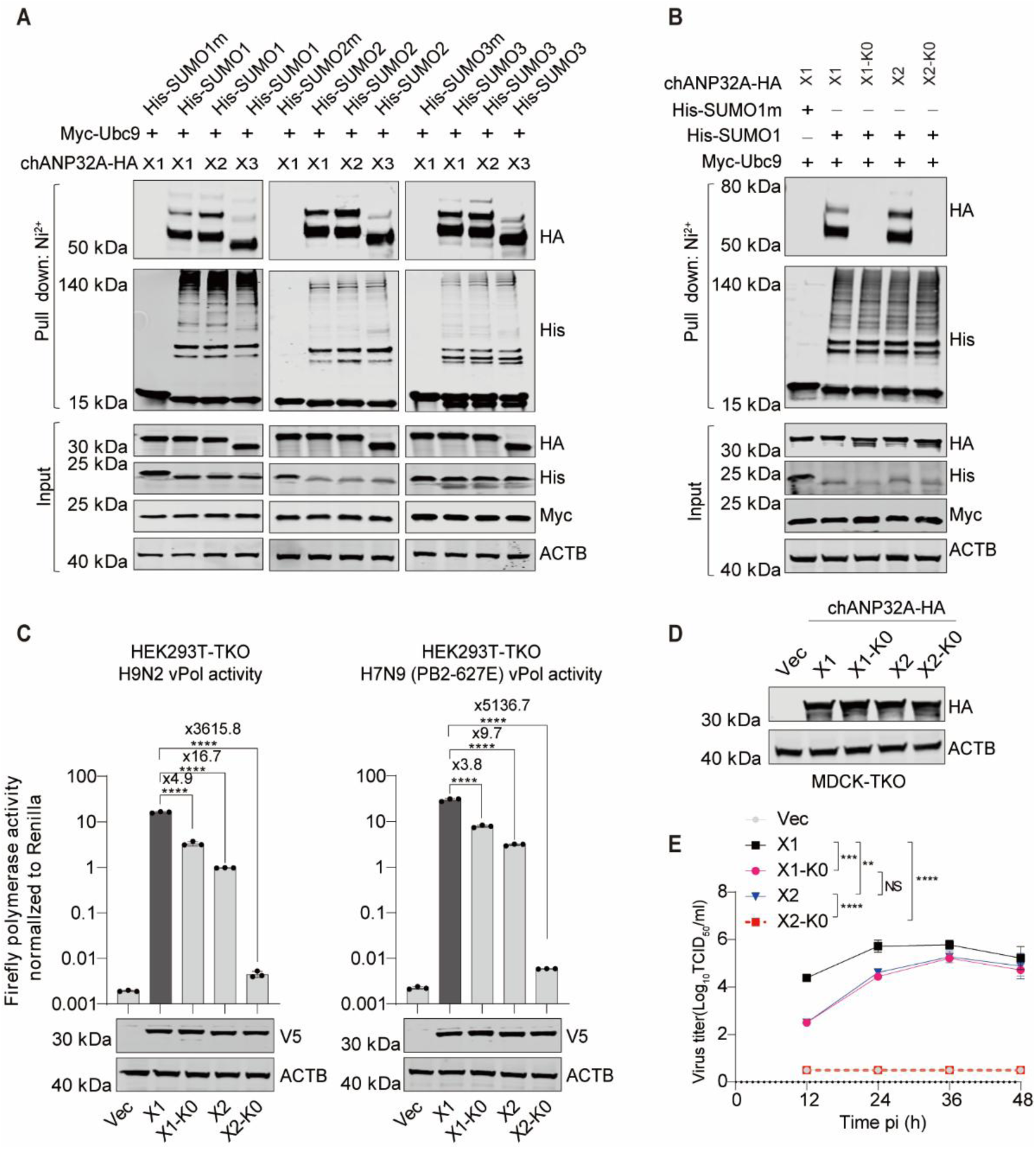
S**UMOylation plays a more essential role in chANP32A-X2 than in chANP32A-X1 for efficient support of AIV polymerase activity (A)** SUMOylation of three chANP32A isoforms by SUMO1, SUMO2, and SUMO3 in HEK293T cells. Lysates from HEK293T cells transfected with the indicated plasmids were subjected to Ni²⁺-NTA bead precipitation under denaturing conditions for SUMOylation assays, followed by immunoblotting analysis. (**B**) SUMOylation was abolished in chANP32A-X1-K0 and chANP32A-X2-K0 mutants in HEK293T cells. (**C**) Polymerase reconstitution assays in HEK293T-TKO cells evaluating the effects of indicated chANP32A-V5 constructs on the polymerase activity of H9N2 and H7N9 (PB2-627E). Statistical analysis and fold changes and relative to X1 are presented. (**D**) Immunoblotting analysis of MDCK-TKO cells transfected with the indicated chANP32A-HA constructs or empty vector. (**E**) Replication kinetics of the avian H9N2 virus in control MDCK-TKO cells or cells transiently expressing X1, X1-K0, X2, or X2-K0 (MOI = 0.001). Viral titers were determined at the indicated time points. In (C) and (E), error bars represent mean ± SD from *n* = 3 independent biological replicates. Statistical significance was determined by one-way ANOVA with Dunnett’s multiple comparisons test (C) or two-way ANOVA (E) (NS, not significant; \*\**p* < 0.01; \*\*\**p* < 0.001; \*\*\*\**p* < 0.0001). In (A) and (B), experiments were independently repeated twice with consistent results.

Taken together, these findings demonstrate that SUMOylation and the SIM act as redundant SUMO-dependent determinants in chANP32A, ensuring robust support of AIV polymerase activity. This compensatory mechanism highlights the evolutionary adaptation of chANP32A to maintain viral fitness despite structural variations.

### SUMOylation at residues K68 and K153 in chANP32A is essential for its role in supporting AIV polymerase activity

To identify critical SUMOylation sites in chANP32A-X2 (which lacks a functional SIM and depends predominantly on SUMOylation), we systematically reverted arginines to lysines in the lysine-free chANP32A-X2-K0 background. Among the 20 lysine residues in chANP32A-X2, reintroduction of K68, K116, or K153 restored H7N9 (PB2-627E) polymerase activity in HEK293T-TKO cells (Fig. 4A and Fig. S2). Double lysine-reverted mutants exhibited synergistic effects, with chANP32A-X2-K0-R68K/R153K demonstrating the strongest support for both H9N2 and H7N9 polymerase activity (Figs. 4B and C). These findings implicate K68 and K153 as essential SUMOylation sites for chANP32A-X2’s role in supporting AIV polymerase activity.

**Figure 4.**
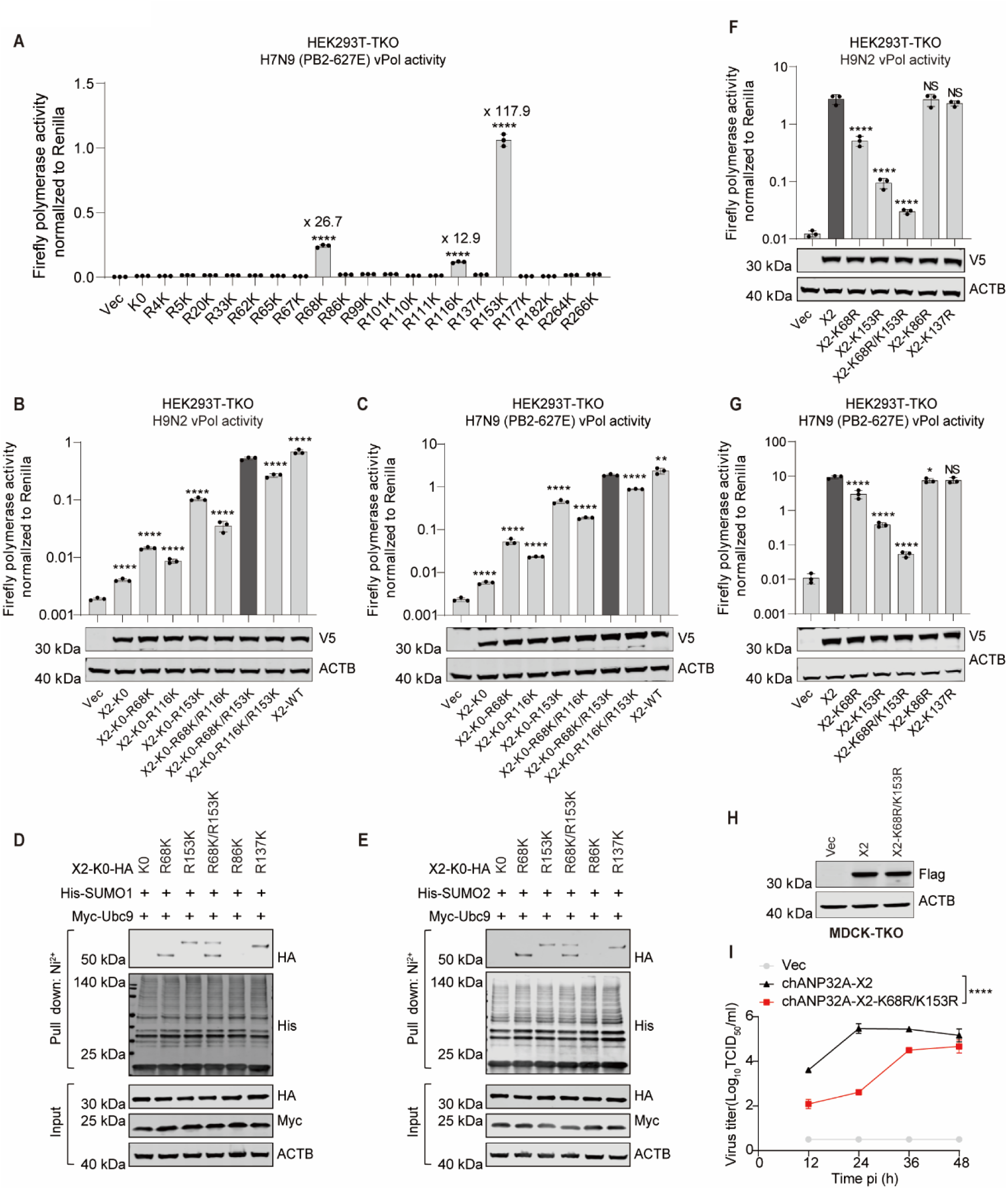
SUMOylation at the K68/K153 sites is essential for chANP32A-X2 function in supporting AIV polymerase activity. **(A)** Polymerase reconstitution assay in HEK293T-TKO cells comparing the impact of the indicated chANP32A-X2-K0 constructs on the activity of the H7N9 (PB2-627E) polymerases. Fold changes and statistical analysis relative to K0 are presented. (**B, C**) Polymerase reconstitution assay in HEK293T-TKO cells evaluating the effects of indicated V5-tagged constructs on the activity of H9N2 (B) and H7N9 (PB2-627E) (C) polymerases. Statistical analysis relative to X2-K0-R68/153K are presented. (**D, E**) Ni²⁺-NTA bead affinity pull-down assays demonstrating SUMO1 (D) or SUMO2 (E) modification at the K68 and K153 sites of chANP32A-X2. (**F, G**) Polymerase reconstitution assay in HEK293T-TKO cells comparing the effect of the indicated chANP32A-X2-V5 constructs on the activity of the H9N2 (F) and H7N9 (PB2-627E) (G) polymerases. Statistical analyses were performed relative to X2. (**H**) Immunoblotting analysis of MDCK-TKO cells stably reconstituted with the indicated chANP32A-X2-Flag constructs or empty vector. (**I**) Replication kinetics of the avian H9N2 virus in control MDCK-TKO cells or cells stably expressing chANP32A-X2 or its mutants (MOI = 0.001). Viral titers were determined at the indicated time points. In (A to C), (F), (G), and (I), error bars represent mean ± SD from *n* = 3 independent biological replicates. Statistical significance was determined by one-way ANOVA with Dunnett’s multiple comparisons (A to C), (F) and (G) or two-way ANOVA (I). (NS, not significant; \**p* < 0.05; \*\**p* < 0.01; \*\*\*\**p* < 0.0001). In (D) and (E), experiments were independently repeated three times with consistent results.

To validate this, SUMOylation assays in HEK293T-SENP1-KO cells revealed distinct ∼50 kDa SUMOylated bands for K68 and K153 mutants, absent in the K0 control (Figs. 4D and E), confirming K68/K153 as SUMOylation sites. For specificity validation, we included non-SUMOylated (K86) and highly SUMOylated (K137) lysines as controls in these experiments. Crucially, the K68R/K153R double mutant displayed markedly reduced AIV polymerase activity compared to wild-type chANP32A-X2 in HEK293T-TKO cells (Figs. 4F and G). Multi-cycle replication assays in MDCK-TKO cells further demonstrated that the K68R/K153R double mutant severely attenuated H9N2 replication compared to wild-type chANP32A-X2 (Figs. 4H and I). Notably, while K137 was SUMOylated, its mutation did not affect polymerase activity (Figs. 4F and G), and global SUMOylation levels remained unchanged in K68R/K153R mutants (Fig. S3), suggesting compensatory SUMOylation at residues like K137 that fail to functionally substitute for K68/K153.

Intriguingly, slower electrophoretic migration of SUMOylated X2-K0-R153K and X2-K0-R137K compared to X2-K0-R68K (Figs. 4D and E) remains unexplained and warrants further study.

Collectively, these findings establish that site-specific SUMOylation at residues K68 and K153—not global SUMOylation—is indispensable for chANP32A-X2 to enable AIV polymerase activity and sustain viral replication.

### The 28-aa insert serves as the third determinant in chANP32A

Like huANP32A/B, chANP32A-X3 lacks the 33 amino-acid insert and exhibits only weak support for AIV polymerase activity (Fig. 1). In contrast, chANP32A-X2 strongly enhances polymerase activity, a functional disparity attributed to its unique 29-amino-acid insert (Fig. 1A). Since chANP32A-X2 requires SUMOylation to promote AIV polymerase activity (Fig. 3 and Fig. 4) and displays slightly higher SUMOylation levels than X3 (Fig. 3A), we hypothesized that its 29-amino-acid insert enhances SUMOylation to boost functionality. The 29 amino-acid insert contains two lysine residues (K177 and K182; Fig. 5A), initially proposed as potential SUMOylation sites. However, mutating these residues (K177R, K182R, or combined) in chANP32A-X2 did not impair its ability to support H9N2 or H7N9 polymerase activity in HEK293T-TKO cells (Figs. 5B and C). In contrast, the K68R/K153R double mutant severely reduced activity below X3 levels, confirming that SUMOylation at K68/K153—not K177/K182—is essential.

**Figure 5.**
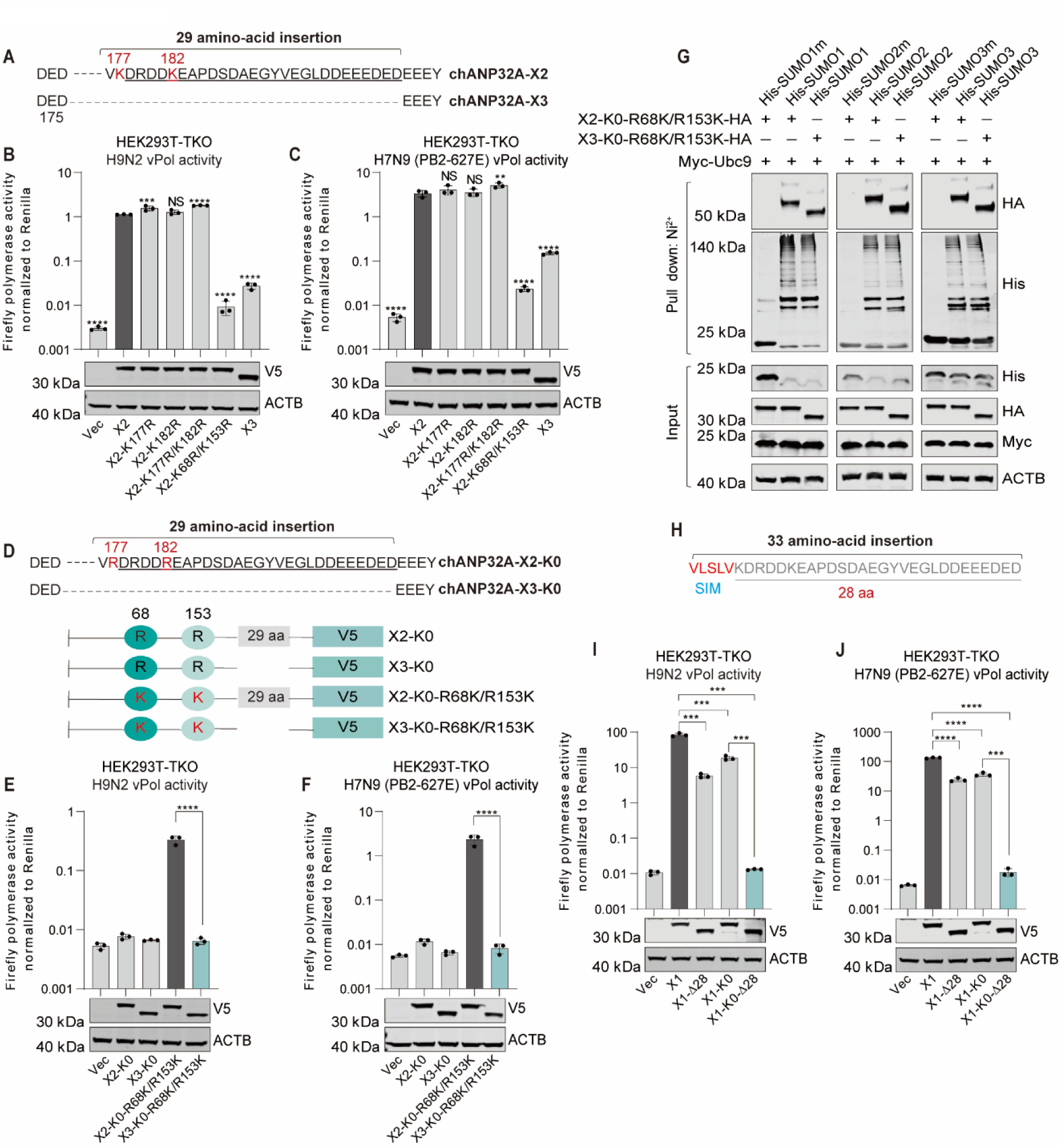
The 28-aa insert is essential for chANP32A to support AIV polymerase activity. **(A)** Sequence alignment of chANP32A-X2 and chANP32A-X3. (**B, C**) Polymerase reconstitution assays in HEK293T-TKO cells evaluating the effect of the indicated V5-tagged constructs on the activity of H9N2 (B) and H7N9 (PB2-627E) (C) polymerases. Statistical analyses were performed relative to X2. (**D**) Diagram depicting the chANP32A-X2-K0-V5 and chANP32A-X3-K0-V5 constructs. (**E, F**) Polymerase reconstitution assay in HEK293T-TKO cells comparing the effect of the indicated constructs on the activity of the H9N2 (E) and H7N9 (PB2-627E) (F) polymerases. (**G**) SUMOylation assays to compare the effect of the 29 amino-acid insert on the SUMOylation levels at K68/K153 of chANP32A. (**H**) Diagram depicting the sequence composition of the 33 amino-acid insert of chANP32A. (**I, J**) Polymerase reconstitution assays in HEK293T-TKO cells evaluating the effect of the indicated V5-tagged constructs on the activity of H9N2 (I) and H7N9 (PB2-627E) (J) polymerases. In (B), (C), (E), (F), (I) and (J), error bars represent mean ± SD from *n* = 3 independent biological replicates. Significance was determined by one-way ANOVA with Dunnett’s multiple comparisons test (B, C, E and F) or unpaired Student’s t-tests (I and J) (NS, not significant; \*\**p* < 0.01; \*\*\**p* < 0.001; \*\*\*\**p* < 0.0001). In (G), experiments were independently repeated three times with consistent results.

To determine whether the 29-amino-acid insert enhances SUMOylation at K68/K153, SUMOylation in chANP32A-X2-K0-R68K/R153K and chANP32A-X3-K0-R68K/R153K was compared. X2-K0-R68K/R153K exhibited stronger polymerase activity than its X3 counterpart (Figs. 5D–F) despite equivalent SUMOylation levels at K68/K153 (Fig. 5G), demonstrating that the insert enhances activity through mechanisms independent of direct SUMOylation.

Notably, the 33-amino-acid insert in wild-type chANP32A (X1) includes a SIM (176-VLSLV-180) and a 28-aa region (Fig. 5H). The X2 isoform retains this 28-aa segment plus V180 from the SIM (Fig. 1A), suggesting the 28-aa region is critical. To test this, we generated chANP32A-X1-Δ28 and chANP32A-X1-K0-Δ28, deleting the 28-aa insert while retaining the SIM. Both mutants exhibited significantly reduced AIV polymerase activity, with chANP32A-X1-K0-Δ28 losing all function (Figs. 5I and J). These results confirm that the 28-aa insert is critical for chANP32A activity but requires SUMOylation to exert its effect.

Collectively, these findings identify the 28-aa insert as a third functional determinant in chANP32A. Its ability to regulate AIV polymerase activity is strictly dependent on SUMOylation, which serves as a prerequisite for its efficacy.

### Synergistic and redundant roles of SIM, SUMOylation, and the 28-aa insert in regulating chANP32A function

Our above findings establish that chANP32A’s ability to support AIV polymerase activity relies on three interdependent functional determinants: the SIM, SUMOylation, and the 28-aa insert. To evaluate whether these determinants act additively or synergistically, we conducted polymerase activity assays in HEK293T-TKO cells expressing chANP32A mutants lacking specific determinants. Wild-type chANP32A (X1) displayed maximal activity, whereas deletion of any single determinant resulted in moderate yet statistically significant functional impairment (Figs. 6A and B). Strikingly, mutants retaining only one determinant (e.g., X3, X2-K0, or X1-K0-Δ28) exhibited uniformly low activity, confirming that no single determinant suffices for robust polymerase support. Conversely, retaining any two determinants restored near-wild-type activity, demonstrating functional redundancy and indicating that two determinants are sufficient for optimal function. Notably, the 28-aa insert function well via either SIM or SUMOylation, highlighting its functional reliance on SUMO-mediated mechanisms.

**Figure 6.**
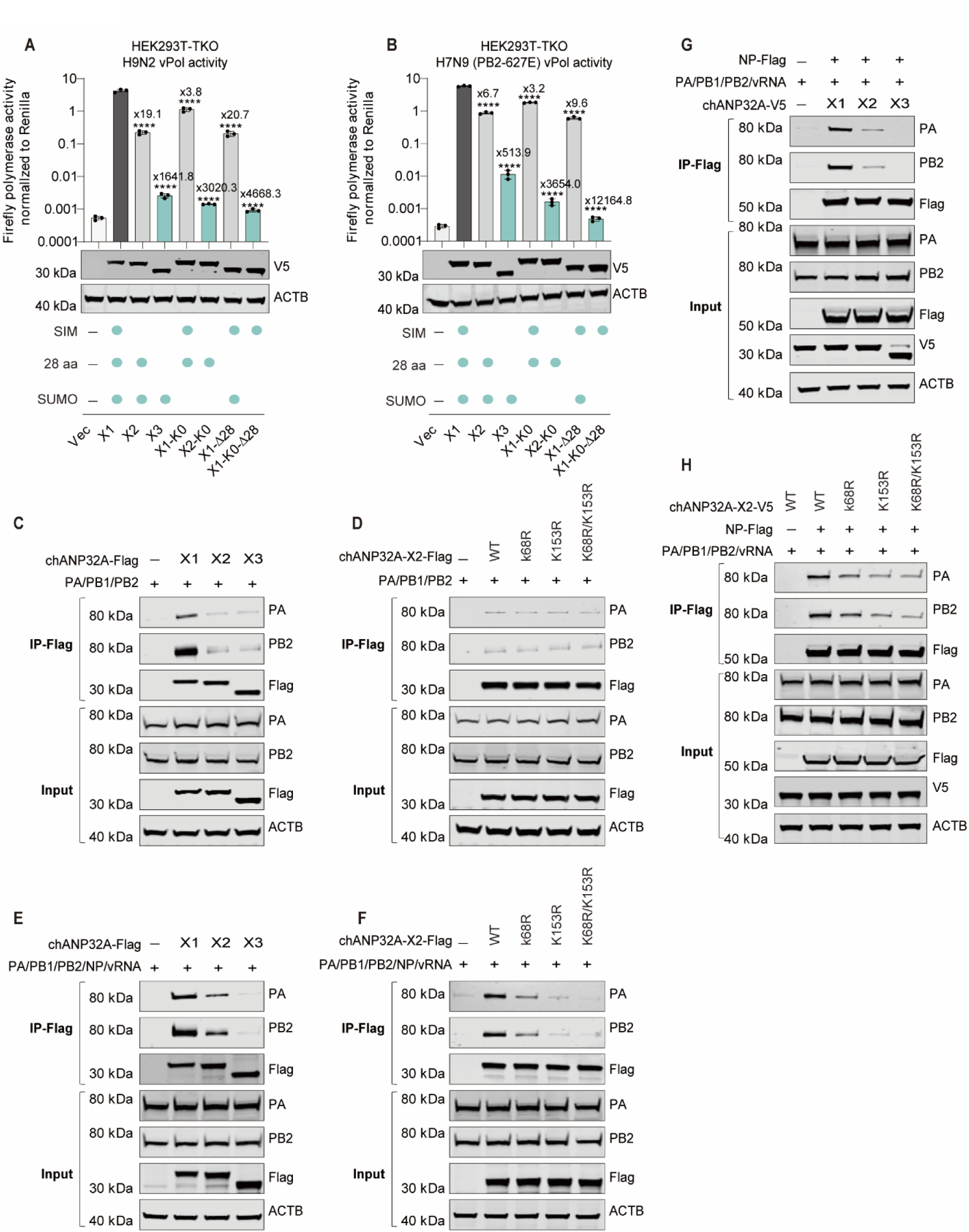
Three determinants in chANP32A exhibit functional redundancy in regulating AIV polymerase activity. (**A**, **B**) Polymerase reconstitution assay in HEK293T-TKO cells revealed that at least two functional determinants are required to support the activity of H9N2 (A) and H7N9 (PB2-627E) (B) polymerases. Error bars represent mean ± SD from *n* = 3 independent biological replicates. Fold changes and statistical analysis relative to X1 are presented. Significance was determined by one-way ANOVA with Dunnett’s multiple comparisons test (\*\*\*\**p* < 0.0001). (**C**) Interaction between the H9N2 polymerase complex and indicated chANP32A isoforms. HEK293T-TKO cells were transfected with chANP32A-Flag constructs (0.4 μg) and plasmids encoding H9N2 polymerase complex (0.2 μg PA, 0.4 μg PB1, 0.4 μg PB2). After anti-Flag immunoprecipitation 24 h post-transfection, proteins were analyzed by western blotting. (**D**) Impact of K68R, K153R, or K68R/K153R mutations on the H9N2 polymerase complex-chANP32A-X2 interaction. Experiments were performed as in Fig. 6C. (**E**) Binding of H9N2 vRNP to indicated chANP32A isoforms. HEK293T-TKO cells were transfected with the indicated chANP32A-Flag constructs (0.4 μg), plasmids encoding H9N2 RNP proteins (0.8 μg NP, 0.2 μg PA, 0.4 μg PB1, and 0.4 μg PB2) together with vRNA-luciferase reporter (0.4 μg). Following anti-Flag precipitation at 24 h post-transfection, the indicated proteins were analyzed with western blotting. (**F**) Effect of K68R, K153R, or K68R/K153R mutations on H9N2 vRNP-chANP32A-X2 interaction. Experiments were performed as in Fig. 6E. (**G**) Analysis of H9N2 vRNP assembly in HEK293T-TKO cells reconstituted with different chANP32A isoforms. Cells were transfected with chANP32A-V5 (0.4 μg), plasmids encoding H9N2 RNP proteins (0.8 μg NP-Flag, 0.2 μg PA, 0.4 μg PB1, 0.4 μg PB2), and the vRNA-luciferase reporter (0.4 μg). Following anti-Flag precipitation at 24 h post-transfection, the indicated proteins were analyzed with western blotting. (**H**) Analysis of H9N2 vRNP assembly in HEK293T-TKO cells reconstituted with chANP32A-X2 and SUMOylation-defective mutants (K68R, K153R, or K68R/K153R). Experiments were performed as in Fig. 6G. In (C-H), experiments were independently repeated at least twice with consistent results.

Prior studies demonstrated that ANP32A/B interacts with IAV vPol to regulate genome synthesis ^20^, while recent work revealed that the AIV NS2 protein enhances AIV vRNP assembly in mammalian cells by strengthening ANP32A/B-vRNP interactions ^36,39^. We hypothesized that the SIM, SUMOylation, and the 28-aa insert in chANP32A collectively enhance its binding affinity for the AIV vPol complex or vRNP, thereby facilitating vRNP assembly. Co-IP assays showed that chANP32A-X1 interacts more robustly with the H9N2 vPol complex than X2 or X3 (Fig. 6C). However, no difference in interaction efficiency was observed between X2 and X3 (Fig. 6C). Similarly, the SUMOylation-deficient X2-K68R/K153R mutant exhibited no altered binding to the vPol complex compared to wild-type X2 (Fig. 6D). In contrast, vRNP interaction assays revealed that chANP32A-X3 poorly co-precipitated H9N2 vRNP components (PB2 and PA), whereas X1 displayed the strongest interaction (Fig. 6E). The K68R/K153R mutation in X2 impaired its ability to bind vRNP (Fig. 6F). Consistent with this, vRNP assembly assays demonstrated that X1 supported the highest levels of PB2/PA-NP co-precipitation, while X3 and SUMOylation-deficient mutants (X2-K68R/K153R) were severely compromised (Figs. 6G and H). These results collectively establish that the SIM, SUMOylation, and 28-aa insert enhance chANP32A’s capacity to interact with and promote AIV vRNP assembly.

Notably, none of these determinants influenced chANP32A’s nuclear localization (Fig. S4).

Taken together, these findings demonstrate that the SIM, SUMOylation, and 28-aa insert in chANP32A act synergistically and redundantly to regulate AIV polymerase activity. At least two determinants are required for optimal function, which is mediated through enhanced AIV vRNP-chANP32A interactions and promotion of AIV vRNP assembly. These findings underscore a cooperative mechanism in which SUMO-dependent processes and structural motifs jointly potentiate chANP32A’s role in supporting AIV polymerase activity.

### Human ANP32A/B rely solely on SUMOylation and therefore have limited capacity to support AIV polymerase activity

Human ANP32A and ANP32B exhibit limited capacity to support AIV polymerase activity due to the absence of the avian-specific 33-amino-acid insert found in chANP32A. This insert contains two discrete functional elements: a SIM and a 28-aa segment. While huANP32A/B undergo SUMOylation^36^—a conserved post-translational modification critical for polymerase support—they lack avian-specific structural elements required for robust AIV activity. To test whether SUMOylation-mediated function is conserved between chANP32A and huANP32A/B, we inserted the SIM, 28-aa segment, or both into huANP32A or its lysine-deficient mutant (huANP32A-K0). In HEK293T-TKO cells, both SIM and 28-aa insert alone restored AIV polymerase activity in huANP32A, but neither rescued SUMOylation-deficient huANP32A-K0 unless combined (Figs. 7A and B), indicating SUMOylation synergizes with either determinant to enable function. In contrast, only the SIM—not the 28-aa insert—conferred activity to huANP32B, and SUMOylation remained essential (Figs. 7C and D). These results confirm that SUMOylation-dependent mechanisms are conserved across species but require species-specific structural elements for full functionality.

**Figure 7.**
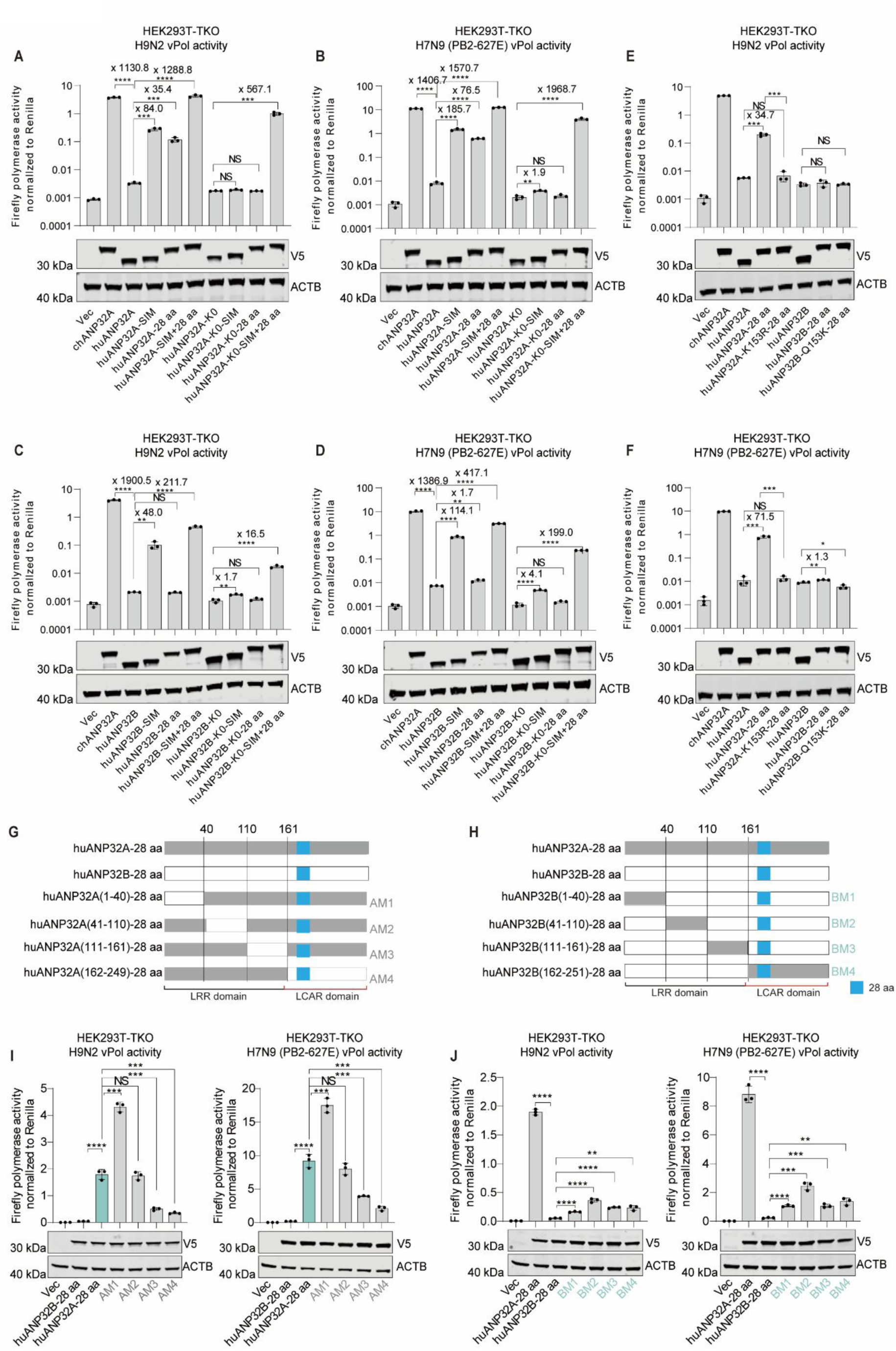
The polymerase supporting activity facilitated by the 28-aa insert in chANP32A is not transferable to huANP32B. (**A, B**) Polymerase reconstitution assays in HEK293T-TKO cells showing that insertion of the SIM or a 28-aa insert rescues the functional defects of huANP32A but not huANP32A-K0 in supporting H9N2 (A) and H7N9 (PB2-627E) (B) polymerase activity. (**C, D**) Polymerase reconstitution assays in HEK293T-TKO cells showing that insertion of the SIM, but not the 28-aa insert, confers on huANP32B a high ability to support H9N2 (C) and H7N9 (PB2-627E) (D) polymerase activity. (**E**, **F**) Polymerase reconstitution assays in HEK293T-TKO cells demonstrating that the non-transferability of the 28-aa to huANP32B is not due to the lack of K153-mediated SUMOylation in huANP32B. Minigenome assays of H9N2 (E) and H7N9 (PB2-627E) (F) polymerase were performed in HEK293T-TKO cells with different ANP32 proteins co-transfected. (**G**, **H**) Schematic diagrams of chimeric clones between huANP32A-28 aa and huANP32B-28 aa, constructed based on known domain architectures. (**I**. **J**) Polymerase reconstitution assays in HEK293T-TKO cells demonstrate that both the LRR and LCAR domains contribute to the functional differences observed between huANP32A-28 aa and huANP32B-28 aa. In (A to F), (I) and (J), error bars represent mean ± SD from *n* = 3 independent biological replicates; Significance was determined by unpaired Student’s t-tests (NS, not significant; \**p* < 0.05; \*\**p* < 0.01; \*\*\**p* < 0.001; \*\*\*\**p* < 0.0001).

The inability of the 28-aa insert to function in huANP32B likely stems from divergent SUMOylation sites: K153, critical in huANP32A, is replaced by glutamine (Q153) in huANP32B. Introducing K153R in huANP32A-28 aa abolished activity, while Q153K in huANP32B-28 aa failed to restore function (Figs. 7E and F), suggesting intrinsic structural differences between huANP32A and huANP32B govern 28-aa compatibility. To pinpoint these differences, we generated chimeric huANP32A/ANP32B-28 aa constructs (Figs. 7G and H). Replacing the LRR (aa 111–161) and LCAR (aa 162–249) domains of huANP32A with huANP32B counterparts abolished 28-aa-mediated activity, whereas swapping analogous regions of huANP32B with huANP32A sequences partially restored function (Figs. 7I and J). This highlights the critical role of the LRR and LCAR domains in enabling 28-aa functionality.

Collectively, these results suggest, while SIM-mediated activity transfers to both huANP32A and huANP32B, the 28-aa insert operates exclusively in huANP32A, dependent on its unique structural contexts. Human ANP32A/B’s reliance solely on SUMOylation—devoid of avian-specific determinants—explains their limited ability to support AIV polymerase, thereby establishing a molecular basis for the species-specific restriction of AIV polymerase in human cells.

## Discussion

Recent structural studies demonstrate that ANP32 proteins act as chaperones, enabling IAV polymerase to form asymmetric dimers critical for genome replication ^24,40,41^. Despite advances, current models remain incomplete in explaining how chANP32A’s 33-residue insert structurally and mechanistically enables selective enhancement of AIV polymerase activity. Here, we identify three functional determinants in chANP32A—the SIM, SUMOylation, and a 28-aa segment within the avian-specific insert—that synergistically and redundantly govern its species-specific support of AIV polymerase.

Previous studies reported minimal impact of SUMOylation on chANP32A-X1 function ^37^, but our work reveals its critical role, likely due to methodological differences: Domingues et al. used HEK293T cells, whereas our ANP32A/B/E triple-knockout HEK293T (TKO) system eliminates endogenous ANP32 interference, enabling precise assessment. We further show that SUMOylation compensates for SIM loss, with both acting via SUMO-dependent mechanisms—non-covalent (SIM) or covalent (SUMOylation)—to regulate chANP32A function. Strikingly, SUMOylation at K68/K153 (but not K137) is essential for chANP32A-X2 activity. However, SUMOylation alone is insufficient in the absence of a functional SIM, as evidenced by chANP32A-X3 retaining only residual polymerase activity. This observation stems from the 28-aa insert acting as a cooperative determinant, which exclusively requires the concurrent presence of SUMO-mediated activities—either through SIM or SUMOylation—to exert its functional role. While both SIM and the 28-aa insert can be transferred to huANP32A, only the SIM confers activity to huANP32B, highlighting intrinsic functional divergences between ANP32 paralogs. These findings resolve how chANP32A’s redundant determinants overcome host restriction and provide a mechanistic framework for AIV adaptation.

Recent studies have identified lysine residues K68/K153 in huANP32A and K68/K116 in huANP32B as bona fide SUMOylation sites^36^. Notably, K68 and K153 are evolutionarily conserved between human and chicken ANP32A. In this work, we further demonstrate that SUMOylation at these conserved residues (K68/K153) is essential for chANP32A’s ability to support AIV polymerase activity, with particularly stringent requirements in the chANP32A-X2 isoform. Intriguingly, structural studies reveal K153 in huANP32A directly engages the PB2 627-domain of the viral polymerase within replication complexes^41^, while K68 in huANP32B maps to the ANP32-polymerase interface^40^, highlighting the functional importance of these residues. While these structural studies depict stabilized interaction states, they may not fully represent dynamic physiological conditions. Our biochemical identification of K68/K153 as SUMOylation sites provides complementary evidence for their functional significance, suggesting post-translational modification could regulate ANP32A-polymerase interactions in a context-dependent manner.

We recently demonstrated that SUMOylation of huANP32A/B facilitates recruitment of the AIV NS2 protein through SUMO-SIM interactions, thereby overcoming species-specific restrictions on AIV polymerase activity in human cells^36^. SUMOylation at K68/K153 is critical for huANP32A-NS2 interaction. Therefore, AIV NS2 could theoretically engage chANP32A via analogous SIM-SUMO interactions. However, our prior work revealed that NS2 minimally impacts chANP32A’s support of AIV polymerase^39^. This apparent paradox is resolved by our findings: SUMOylation in chANP32A operates synergistically with either the SIM or the 28-aa insert to sustain polymerase activity. These redundant mechanisms—SUMO-dependent pathways (covalent SUMOylation and SIM interactions) and a SUMO-independent structural motif (28-aa insert)—render NS2’s contribution functionally redundant in avian cells. Conversely, huANP32A/B, lacking the SIM and 28-aa insert, rely solely on SUMOylation, which is insufficient for robust polymerase activity unless potentiated by NS2. This evolutionary divergence underscores how chANP32A’s multi-layered redundancy ensures robust viral support, while huANP32A/B’s limited adaptability explains their dependence on NS2-mediated enhancement. These insights illuminate how modular domain adaptations in ANP32A/B balance redundancy and constraint across species, shaping host-specific viral fitness.

The IAV polymerase operates within vRNP complexes to drive viral genome replication and transcription ^42^. Restriction of AIV polymerase activity in mammalian cells has been attributed to defective AIV vRNP assembly^6^, with assembly efficiency markedly reduced in the presence of huANP32A/B but not chANP32A^39^. We recently demonstrated that the AIV NS2 protein enhances avian polymerase efficiency by promoting AIV vRNP-ANP32 interactions and AIV vRNP assembly in mammalian systems expressing huANP32A/B, but not in those with chANP32A^39^. Here, we reveal that chANP32A’s ability to support AIV polymerase activity—mediated by three functional determinants (the SIM, SUMOylation, and the 28-aa insert)—is also directly linked to its capacity to strengthen AIV vRNP-chANP32A interactions and enhance vRNP assembly.

In summary, we demonstrate that chANP32A utilizes three functionally redundant yet mechanistically interdependent determinants—a SIM, SUMOylation modifications, and a 28-aa insert—to cooperatively enhance AIV polymerase activity. This enhancement arises from strengthened AIV vRNP-chANP32A binding and facilitated AIV vRNP assembly. In contrast, huANP32A/B, which lack both the SIM and the 28-aa insert, rely solely on SUMOylation and fail to sustain comparable polymerase support, thereby establishing a structural and evolutionary barrier to AIV cross-species adaptation. These findings reveal how modular domain plasticity in ANP32A/B proteins balances functional redundancy with species-specific constraints, driving host-adapted viral fitness. By elucidating these mechanisms, our work provides a molecular framework for targeting ANP32A/B adaptability in antiviral strategies aimed at disrupting host-pathogen coevolution.

## Materials and Methods

### Cells, plasmids construction and chemical reagents

HEK293T cells, *ANP32A/ANP32B/ANP32E* triple-knockout HEK293T cells (HEK293T-TKO)^20^, *SENP1*-knockout HEK293T cells (HEK293T-SENP1-KO)^43^, MDCK cells, and *ANP32A/ANP32B/ANP32E* triple-knockout MDCK cells (MDCK-TKO)^43^ were cultured in DMEM supplemented with 10% fetal bovine serum (Alphabio, Tianjin Alpha Biotechnology Co., Ltd) and 1% penicillin-streptomycin (Gibco). Cells were maintained at 37°C in a 5% CO_2_ atmosphere. Transfection experiments were performed using polyethyleneimine (PEI) reagent.

Plasmids used in this study, including VR1012-Flag-human SENP1/2/3/5/6/7, VR1012-Flag-human USPL1, VR1012-human PIAS1/PIAS2α/PIAS2β/PIAS3/PIAS4, pCAGGS-Myc-Ubc9, pCEF-His-SUMO1/2/3, and pCEF-His-SUMO1m/2m/3m (the latter containing GG-to-AA mutations at the C-terminus of His-tagged SUMO1/2/3 essential for SUMO conjugation), were previously described^39,43^. Plasmids pCAGGS-chANP32A-X1/X2/X3-Flag/V5 and VR1012-chANP32A-X1/X2/X3-HA, along with their various mutants, were generated using standard molecular cloning techniques. Gene sequences for the lysine-free mutants of chANP32A-X1 and chANP32A-X2 were synthesized and inserted into either the pCAGGS or VR1012 expression vector. Additional gene variants and specific mutations were introduced using PCR-based methods and confirmed by DNA sequencing. Reporter constructs and pCAGGS expression plasmids encoding polymerase components and NP for A/chicken/Zhejiang/B2013/2012 (H9N2) and A/Anhui/01/2013 (H7N9) were previously described^20,26^. All constructs were validated by sequencing. Lentiviral stocks were generated using pLVSIN-PGK-Puro-cloned variants.

Puromycin (Catalog #A1113803) was acquired from Thermo Fisher Scientific, while subasumstat (TAK-981; HY-111789), a small-molecule inhibitor of the SUMOylation enzymatic cascade, was sourced from MedChemExpress

### Establishment of stable cell lines

MDCK-TKO cells stably expressing Flag-tagged chANP32A variants were generated via lentiviral transduction (pLVSIN-CMV-PGK-puro vector). Puromycin (1 μg/ml) selection began 48 hours post-transduction.

### Virus stock production, and infection assays

Virus rescue was performed in HEK293T cells by co-transfecting the H9N2 reverse genetics plasmids and chANP32A-Flag constructs. Supernatants were harvested at 48 hpi and propagated in SPF chicken embryos (35°C, 48 h). Viral titers (TCID₅₀) were determined in MDCK-chANP32A (X1) cells^44^.

MDCK-TKO cells reconstituted with chANP32A variants were infected with H9N2 (MOI = 0.001) in opti-MEM. Post-adsorption (1–2 h), medium was replaced with opti-MEM + 1 μg/ml TPCK-trypsin. Supernatants were collected at indicated time points for TCID₅₀ titration.

### Immunoprecipitation assays

Lysates from transfected cells were prepared using lysis buffer (50 mM Hepes-NaOH [pH 7.9], 100 mM NaCl, 50 mM KCl, 0.25% NP-40, and 1 mM DTT) supplemented with a complete inhibitor cocktail (APExBIO, Houston, USA; K1007). After centrifugation at 12,000 rpm for 10 minutes, the clarified supernatants were incubated overnight at 4°C with anti-Flag M2 magnetic beads (Sigma-Aldrich, M8823). The beads were then washed three times with lysis buffer, and bound proteins were eluted using 3×Flag peptide (APExBIO, Houston, USA; A6001). Immunoprecipitated proteins were subsequently analyzed by western blotting.

### SUMOylation assays

Transfected cell lysates were prepared using RIPA lysis buffer supplemented with 1% SDS, 10 mM NEM, and a complete inhibitor cocktail (APExBIO, Houston, USA; K1007). After sonication to ensure homogeneity, the lysates were centrifuged at 12,000 rpm for 10 minutes. The clarified supernatants were incubated overnight at 4°C with Ni2+-NTA beads (Sangon Biotech, C650033). The beads were then sequentially washed with buffer A (50 mM Tris-HCl, 0.5 M NaCl, 6 M guanidinium-HCl), buffer B (50 mM Tris-HCl, 0.5 M NaCl, 8 M urea), and PBS. Finally, bead-bound proteins were eluted using elution buffer containing 200 mM imidazole and analyzed by western blotting.

### Western blots

Western blot analysis was conducted following standard procedures as described previously^20^. The antibodies utilized in this study included: rabbit anti-Flag (Sigma-Aldrich, F7425), rabbit anti-HA (Sigma-Aldrich, H6908), rabbit anti-ACTB (Abclonal, AC026), mouse anti-ACTB (Abclonal, AC004), rabbit anti-Myc (Abclonal, AE070), mouse anti-His (Proteintech, 66005-1-Ig), rabbit anti-V5 (Proteintech, 14440-1-AP), rabbit anti-SENP1 (Proteintech, 25349-1-AP), rabbit anti-PIAS2 (Proteintech, 16074-1-AP), rabbit anti-influenza A virus PB2 (GeneTex, GTX125926), rabbit anti-influenza A virus PA (GeneTex, GTX118991), mouse anti-influenza A virus PA (produced in our laboratory, 1:5000 for WB), DyLight 800 labeled Anti-Mouse IgG (H+L) Antibody (KPL, 5230-0415) and DyLight 680-labeled Anti-Rabbit IgG (H+L) Antibody (KPL, 5230-0402).

### Minigenome assays

HEK293T-TKO cells were seeded into 24-well plates and co-transfected with the following plasmids: pPolI-luc (40 ng), *Renilla* luciferase expression plasmid (5 ng), pCAGGS-PB1 (20 ng), pCAGGS-PB2 (20 ng), pCAGGS-PA (10 ng), pCAGGS–NP (40 ng), and plasmids encoding various ANP32 genes (20 ng each). Cells were lysed 24 hours post-transfection, and luciferase activity was assessed using the Dual-Glo Luciferase Assay System (Promega) with a Berthold Centro LB 960 microplate luminometer.

### siRNA treatment

siRNA transfection was carried out using Lipofectamine™ 2000 according to the manufacturer’s protocol on the same day HEK293T cells were seeded. Twenty-four hours post-siRNA transfection, cells underwent a second transfection with indicated plasmids using PEI transfection reagent. Cells were collected for analysis 24 hours after this second transfection step. Cells were harvested for analysis 24 hours after the second transfection step. Non-targeting control and siRNA oligonucleotides targeting SENP1 and PIAS2, previously described^43^, were obtained from Seven Innovation (China, Beijing) Biotechnology.

### Immunofluorescence

The specified cells were fixed, permeabilized, and stained with primary and secondary antibodies following a previously described protocol^45^. Images were acquired using a confocal microscope (Carl Zeiss LSM 800) and analyzed with ZEN 2.3 LITE software.

### Statistical analysis

Quantitative data are expressed as mean ± SD. Statistical differences were analyzed using one-way ANOVA with Dunnett’s multiple comparisons test, unpaired Student’s t-test, or two-way ANOVA, with GraphPad Prism 7.0 software. Statistical parameters are reported in the figures and figure legend**s.**

## Acknowledgments

We are grateful to Professor Hualan Chen (Harbin Veterinary Research Institute, CAAS) for providing plasmids and for her insightful discussions. We also thank Dr. Zejun Li (Shanghai Veterinary Research Institute, CAAS) for providing plasmids. This work was supported by the National Natural Science Foundation of China (grants 32330103 and 32302959)

## Author Contributions

X.J.W. supervised the study and revised the manuscript; X.J.W. and L.K.S. designed the study and analyzed the data; L.K.S. wrote the original draft and performed the experiments; M.M.Y. and Y.X.Q. assisted in sample collection; X-F.W. contributed to manuscript editing. All authors reviewed and approved the final manuscript.

## Supplementary figures

**Fig. S1.**
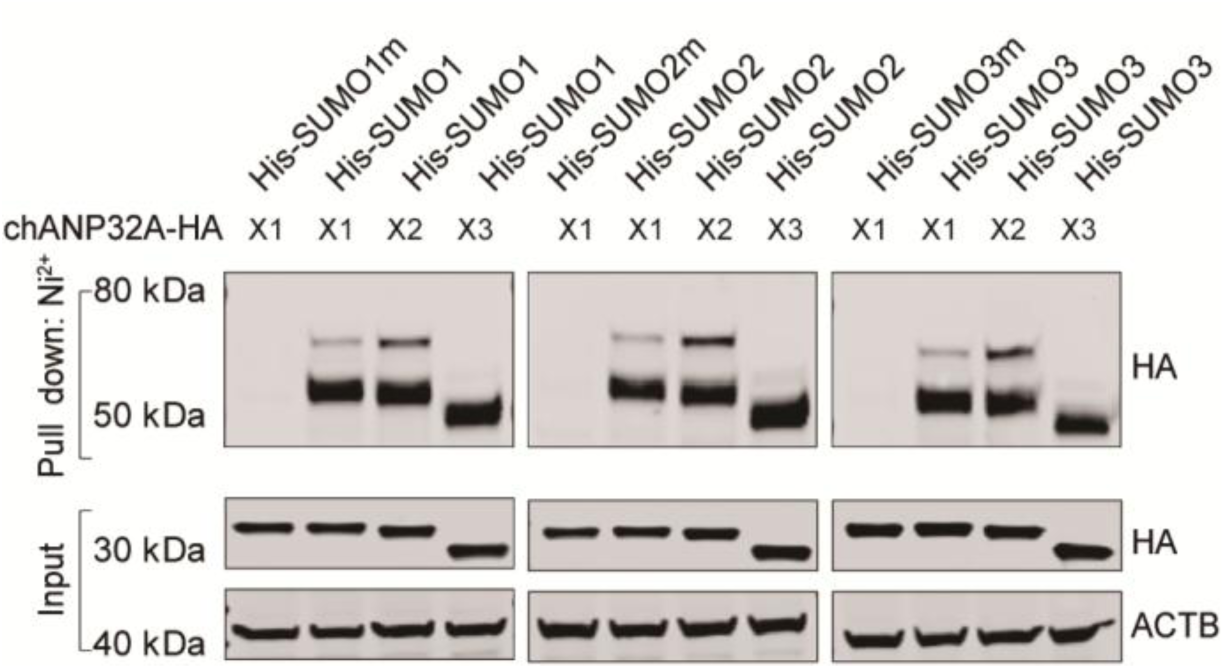
The three chANP32A isoforms exhibit a similar SUMOylation pattern. Lysates from HEK293T cells transfected with the indicated plasmids were subjected to Ni²⁺ -NTA bead precipitation under denaturing conditions for SUMOylation assays, followed by immunoblotting analysis. This experiment was independently repeated three times with consistent results.

**Fig. S2.**
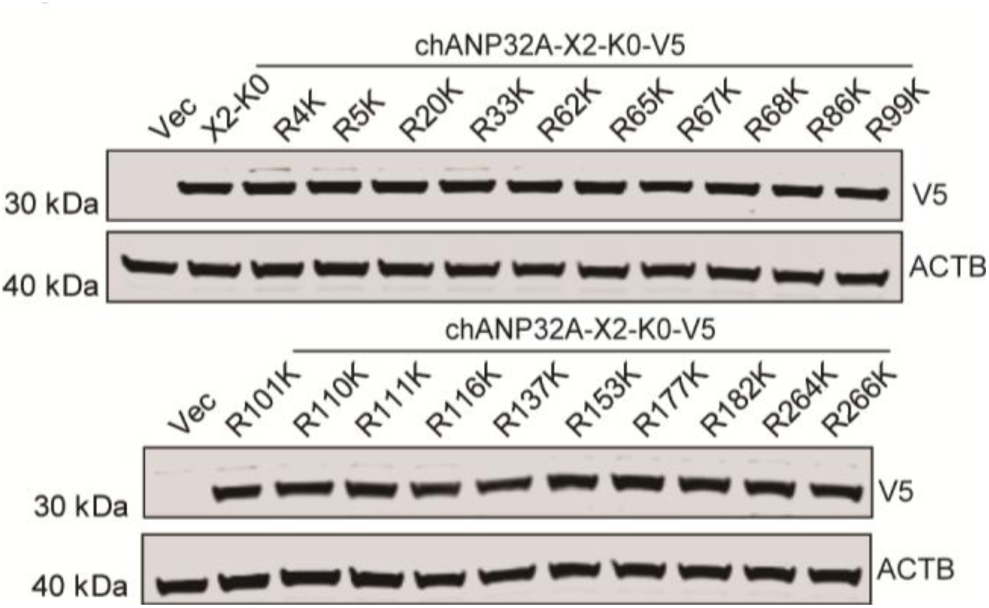
Immunoblotting analysis of lysates from HEK293T-TKO cells transfected with the indicated constructs. HEK293T-TKO cells were transfected with the specified plasmids for 24 hours, after which they were harvested for immunoblotting analysis using specific antibodies.

**Fig. S3.**
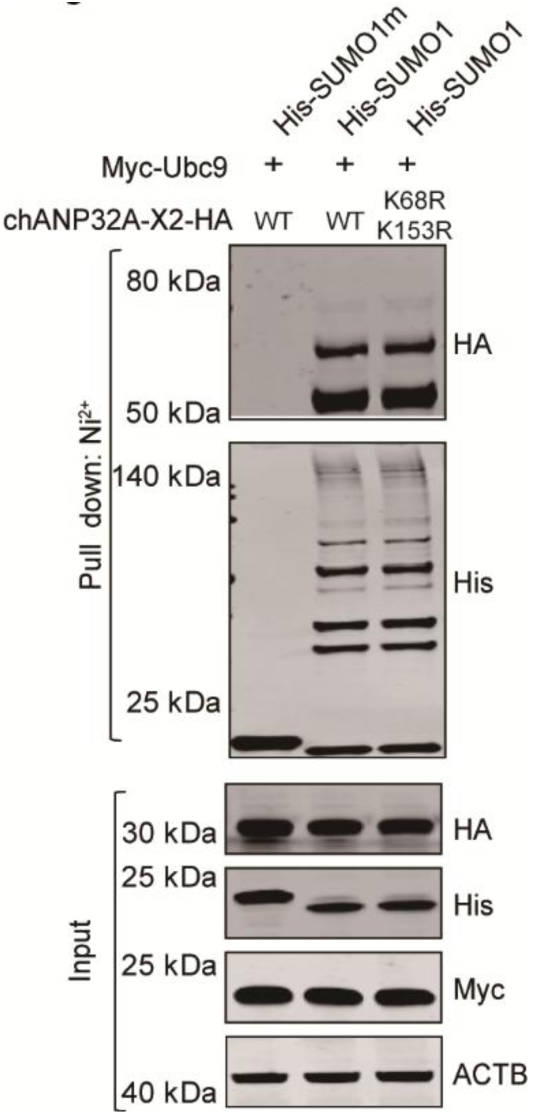
K68R/K153R mutations in chANP32A-X2 do not affect its overall SUMOylation levels. HEK293T cells were transfected with His-SUMO1 or its non-conjugatable form, His-SUMO1m, along with Myc-Ubc9 and either chANP32A-X2-HA or its mutants for 24 hours. SUMOylation assays and immunoblotting analysis were then performed.

**Fig. S4.**
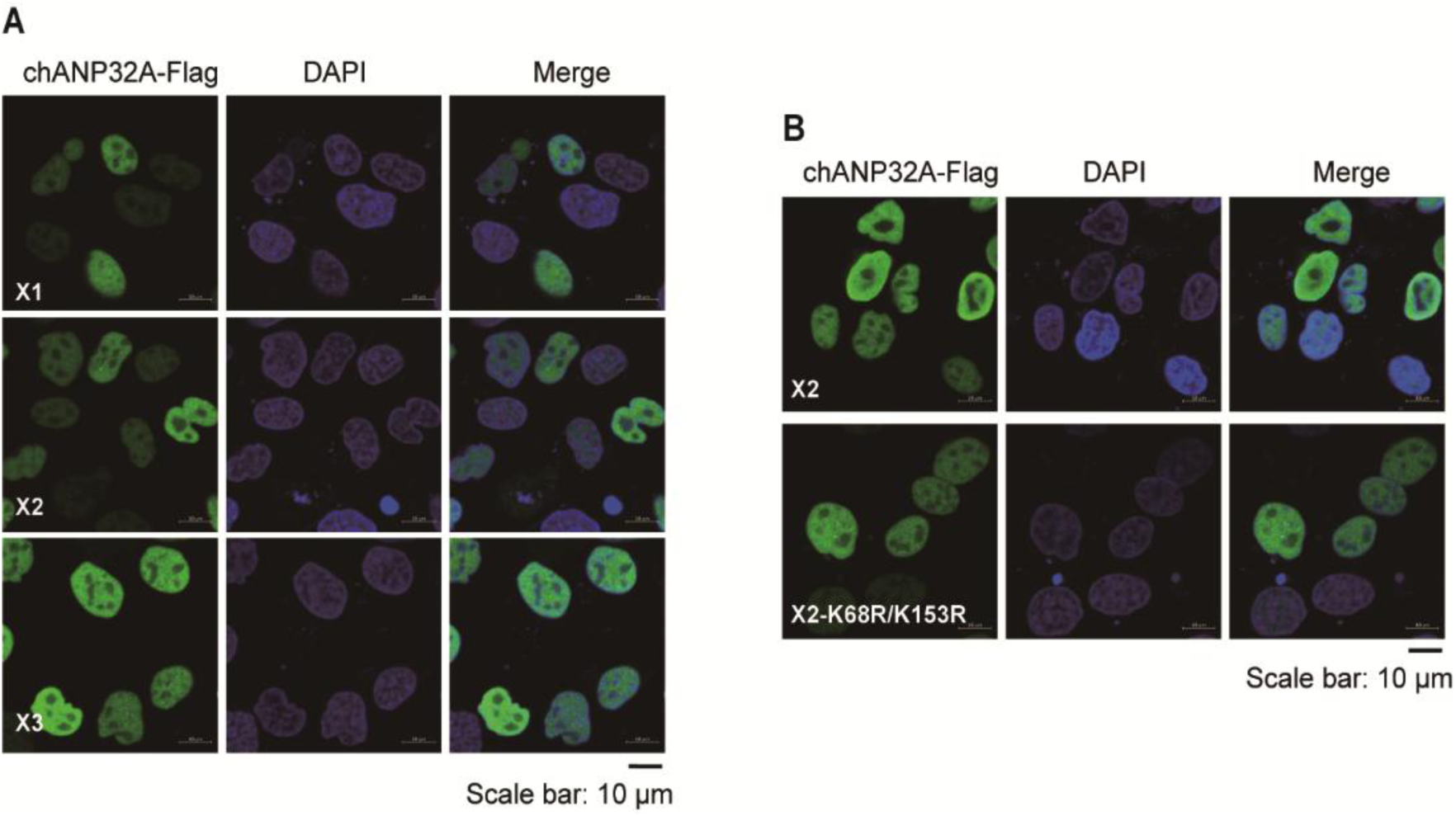
Three determinants within chANP32A do not affect its nuclear localization. **(A)** MDCK-TKO cells stably expressing chANP32A-X1, chANP32A-X2, or chANP32A-X3 were analyzed by immunofluorescence. (**B**) MDCK-TKO cells stably expressing chANP32A-X2 or its mutant were analyzed by immunofluorescence.

